# Annual changes in the Biodiversity Intactness Index in tropical and subtropical forest biomes, 2001-2012

**DOI:** 10.1101/311688

**Authors:** Adriana De Palma, Andrew Hoskins, Ricardo E. Gonzalez, Luca Börger, Tim Newbold, Katia Sanchez-Ortiz, Simon Ferrier, Andy Purvis

## Abstract

Few biodiversity indicators are available that reflect the state of broad-sense biodiversity—rather than of particular taxa—at fine spatial and temporal resolution. The Biodiversity Intactness Index (BII) estimates how the average abundance of native terrestrial species in a region compares with their abundances before pronounced human impacts. BII is designed for use with data from a wide range of taxa and functional groups and for estimation at any resolution for which data on land use and related pressures are available. For each year from 2001 to 2012, we combined models of how land use and related pressures in tropical and subtropical forested biomes affect overall abundance and compositional similarity of plants, fungi, invertebrates and vertebrates, with data on anthropogenic pressures to produce annual maps of modelled BII at a spatial resolution of 30 arc seconds (roughly 1 km at the equator) across tropical and subtropical forested biomes. This is the first time temporal change in BII has been estimated across such a large region. The approach we have used to model compositional similarity uses data more efficiently than that used previously when estimating BII. Across tropical and subtropical biomes, BII fell by an average of 1.9 percentage points between 2001 and 2012, with 81 countries seeing an average reduction and 43 an average increase; the extent of primary forest fell by 3.9% over the same period. Changes are not strongly related to countries’ rates of economic growth over the same period.

## Introduction

Biodiversity indicators can play an essential role in tracking progress towards policy targets, especially if the indicators link strongly to both the targets and biodiversity, have broad geographic coverage, and are available as a time series [1]. These stringent criteria have contributed to a strong taxonomic bias in biodiversity indicators [1–4]. In an assessment of whether the rate of biodiversity loss had fallen by 2010 [4], only one of four measures of the state of biodiversity considered any non-vertebrate data (the Red List Index considered corals in addition to birds, mammals and amphibians) and none of the three indicators of benefits accrued from biodiversity did so. This bias is, if anything, stronger among indicators considered in a mid-term analysis of progress towards the Aichi 2020 Targets [1]: only one of the nine measures of the state of biodiversity (coral reef cover) considered non-vertebrate data, and none of the three measures of benefits did so. Indicators based on a taxonomically-broad sets of species are urgently needed [5] because species in different clades often respond differently to given pressures [6–9].

In addition to taxonomic bias, available biodiversity data show strong geographical biases. Many tropical regions tend to be under-represented relative to well-recorded regions such as North America, Western Europe and Australia [10–12], for a range of socioeconomic reasons [13]. If locations are biased relative to the distribution of anthropogenic pressures, the average trend inferred from the data may not accurately reflect the true global trend [14, 15] unless the bias is corrected [e.g. 10].

The Biodiversity Intactness Index [BII: 16] is designed to reflect the status of a much broader set of taxa than most other indicators. BII is defined as ‘the average abundance of a large and diverse set of organisms in a given geographical area, relative to their reference populations’ [16]. The reference condition is approximated as the contemporary situation in minimally-impacted sites, given the paucity of sufficiently precise historical baseline data [16]. BII was estimated originally using expert judgement in lieu of detailed primary biodiversity data [16]. More recently, BII was estimated globally for the year 2005 by combining global maps of pressures with two statistical models (one of the effects of land use and related pressures on overall sampled organismal abundance; one of how land use affects compositional differences between disturbed habitats and primary vegetation [17]. Both models are described in more detail by [15]). These statistical models were fitted to a global compilation of studies, within each of which a comparable ecological survey had been undertaken at multiple sites facing different pressures [described in 18], with a total of 39,123 plant, fungal or animal species and 18,659 sites spanning all 14 of the world’s terrestrial biomes [17]. The models assumed that the relationships between pressures and biodiversity do not vary regionally, but the underlying database [19] has now grown to the point that it can support some region- or biome-specific modelling. This possibility is important given the likelihood of regional variation in responses [e.g. 20–23].

The global estimates for BII in the year 2005 made use of fine-scale (30 arc-second, approximately 1km-resolution) maps of land use [24] produced through statistical downscaling of global 0.5-degree harmonized land-use data [25]. Remote sensing has greatly improved the ability to track land-cover change at fine spatial and temporal resolutions [26]. Methods such as those developed in [24] can in principle convert such data to time-varying land-use estimates, providing a time series of pressure data that can be used to estimate BII annually, greatly enhancing its usefulness as an indicator. We focus here on forest biomes given that detection of forest loss by remote sensing is better developed than detection of grassland loss [27].

Tropical and subtropical forest biomes cover about 17% of the globe’s land surface but are home to most of the world’s terrestrial species, and the ecosystem services they provide sustain well over 1 billion people [28]. Land-use change is the major threat globally to tropical forest biodiversity [29] and is driven by a combination of factors that include agricultural expansion, timber extraction and infrastructure development [30], though rates and patterns of forest loss differ regionally [28]. Deforestation and degradation reduce local species richness across a range of groups [9, 20, 31] but no biodiversity indicators are yet available that give a taxonomically broad picture of the status and trend of tropical forest biodiversity that is well resolved spatially and temporally. Here we use annual global fine-resolution maps of land use and human population density to map modelled BII across the world’s tropical and subtropical forests for each year from 2001 to 2012. Summary statistics of average change in BII at national and regional levels (see Appendix Table 1) will be relevant for biodiversity assessments such as those being undertaken by the Intergovernmental Science-Policy Platform on Biodiversity and Ecosystem Services (IPBES) and the Group on Earth Observations (GEO). We also explore how per capita Gross Domestic Product (GDP) is related to average change in BII across countries. The relationship between economic indicators and biodiversity is still unclear. Recent evidence has suggested that increases in GDP are correlated with increased forest area in tropical forested regions [32], while forest degradation and loss is concentrated in poorer countries [33]; however, other work has shown that increased GDP is correlated with increased forest change [34] and extinction risk in mammals [35].

## Results

BII is estimated by combining statistical models of how land use and related pressures affect both overall organismal abundance and compositional similarity to a minimally-impacted primary forest baseline. Total abundance was significantly influenced by interactions between land use and both human population density (*χ*^2^ = 22.23, df = 10, *p <*0.05) and road density at the 50km^2^ scale (*χ*^2^ = 25.10, df = 10, *p <*0.01). For the model of compositional similarity, all terms were maintained during model simplification (each significant at the *p <*0.01 level according to the permuted likelihood ratio tests).

On average across the tropical and subtropical forested biomes, BII fell from 63.6% in 2001 to 61.7% in 2012 (Fig 1), a rate of loss of approximately 0.17% per year. Losses in BII were most severe in the Tropical and Subtropical Moist Broadleaf Forests (with a loss of 2.03 percentage points). All regions saw an average decline in BII over the period, with Asia and the Pacific suffering the greatest losses (−2.3 percentage points) and the least severe declines in Africa (−1.7 percentage points); similarly, Asia and the Pacific suffered the sharpest declines in the area of primary vegetation (−6.0% relative to the area in 2001), while Africa saw a minor average increase (+3.1%).

**Figure 1.**
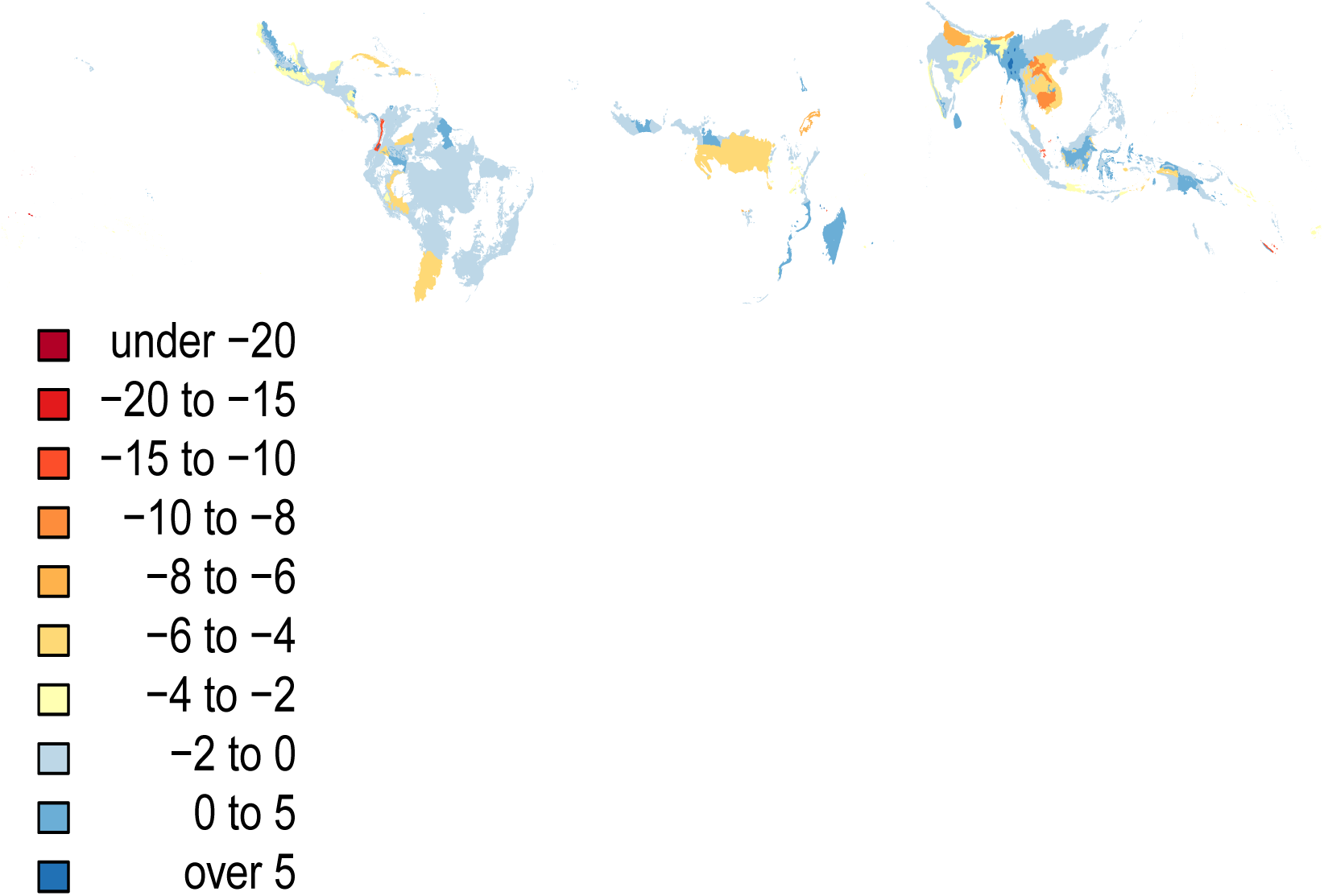
Map of country level differences in BII between 2001 and 2012 (expressed as percentage point difference).

Changes in BII from 2001 to 2012 varied among countries (Fig 2), but the median log response ratio was significantly negative (median = −0.01, Wilcoxon test: V = 2814, p-value *<*0.01). When considering only those countries where at least 50% of their area is within the included tropical or subtropical forest biomes, most countries showed average losses over the time period (Fig 2). Average change over time at the country level was not clearly related to changes in gross domestic product per capita (Fig 3 and Fig S1).

**Figure 2.**
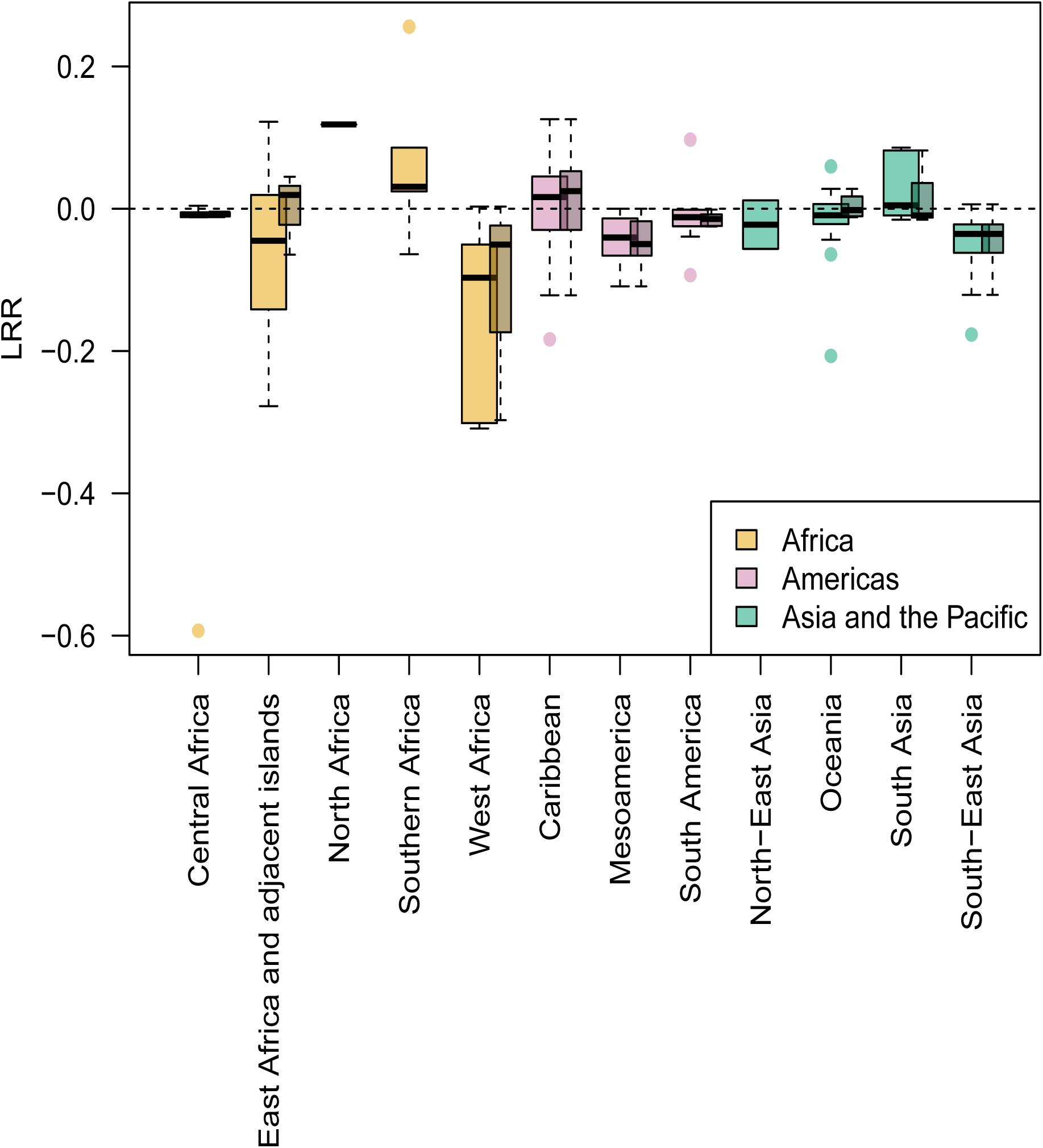
Average change in BII over time at the country level, across different subregions. Change was calculated as the log-response ratio (LRR) of 2012 and 2001 values. A value of zero indicates no change (identified by the dashed line), negative values indicate a decline over time, and positive values indicate an increase in BII over time. Wider, lighter boxes include all countries; narrower, darker boxes use data for countries where BII has been calculated for at least 50% of their area. The center line of the boxplot indicates the median value, boxes show data within the 25th to 75th percentiles, whiskers show points that are up to 1.5 × the interquartile range of the data, points are data that fall outside of these limits.

**Figure 3.**
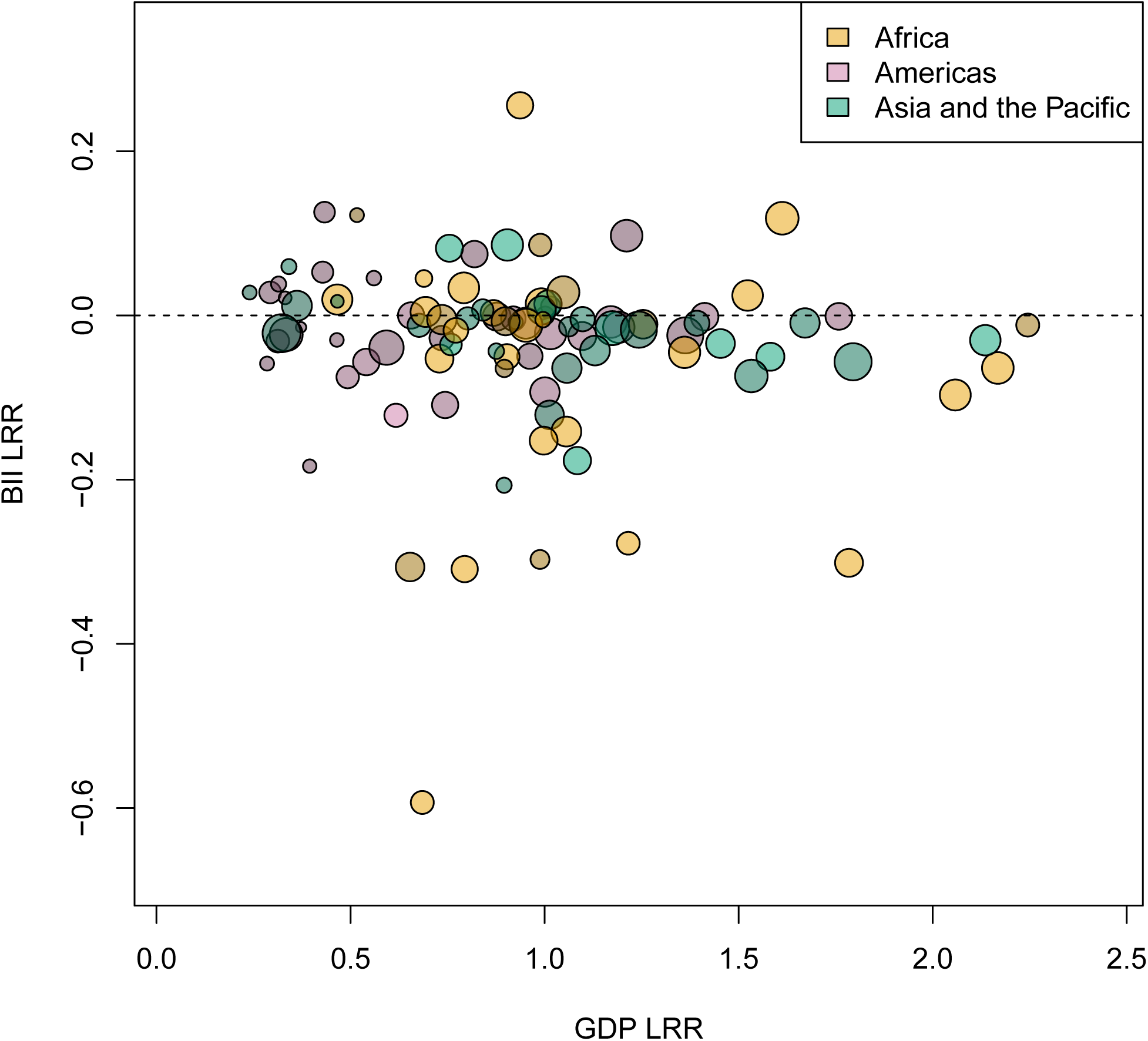
Change in BII over time plotted against the change in GDP per capita for each country. Change was calculated as the log-response ratio of 2012 and 2001 values. A value of zero indicates no change, negative values indicate a decline from 2001 to 2012, and positive values indicate an increase in BII between 2001 and 2012. Note that not all countries have available data on GDP per capita for the years 2001 and 2012 so some countries are excluded from this plot. Colours represent the different regions; colours are also shaded light to dark to represent low, middle and high starting values of GDP. The circles are scaled according to the country area (log10-transformed).

## Discussion

Although human impacts on the rate of global species extinction have perhaps attracted more concern [36–38], local diversity matters more than global diversity for reliable provision of many ecological functions and services. Reduction in local diversity is associated with reduced rates of delivery of key functions [39] as well as greater variance in those rates [40]. This closer link to function is one reason why the Biodiversity Intactness Index (BII) was proposed as a metric that could be used to assess the state of biodiversity relative to its proposed ‘Planetary Boundary’ [41, 42]. In addition, losses across trophic groups can have larger impacts on ecosystem function than losses within a trophic group [40] so, by including multiple taxa, BII may be more functionally relevant than many other measures of local diversity. BII usefully complements indicators focusing on species populations or extinction [43].

BII is likely to be a useful indicator of the state of biodiversity in tropical forests, where single taxonomic groups can be inefficient surrogate indicators for the responses or diversity of other groups [6, 44, although see 45]. In addition, the computation of BII used here combines both alpha diversity (total abundance) and beta diversity (compositional similarity) to estimate the average local abundance of ‘originally-present species’. These two aspects of diversity can show contrasting patterns [46] and responses to human impacts [47, 48]. The contrast may in part help to resolve the recent debate on how human impacts have been affecting the diversity of local ecological assemblages [49].

Steffen et al. [42] proposed a global ‘safe limit’ for biosphere integrity at a BII value of 90%, with a lower bound of 30%. It is unclear whether such a threshold truly exists either globally or across particular biomes as analysed here, or what value it takes if so [41, 50]. However, our results suggest that BII in the world’s tropical forests had already fallen far below the proposed upper bound by 2001 with an average value of 63.6%. Such a low value is unsurprising, given that the highest rates of deforestation in tropical forests occurred during the 1980s and 1990s [51].

Within-country trends over the time period showed substantial variation, but most countries (and all but three subregions) showed declines in average BII. Only three countries are estimated to have remained above the 90% BII threshold: French Guiana, Suriname and Papua New Guinea. This provides a worrying picture for the state of biodiversity in tropical and subtropical forest biomes. Land-use conversion was of course an important predictor of biodiversity loss, but degradation at local (1km) and broader (50km) scales were also significant contributors: for primary vegetation, some of the strongest declines in compositional similarity were seen as road density increased at broader spatial scales, particularly in lightly and intensively-used primary vegetation. This finding suggests that natural intact vegetation must be protected at varying spatial scales in order to conserve local diversity [52]. Indeed, the amount of natural intact vegetation is low and continuing to decline [53–55]. However, despite the suggestion that forest loss and degradation tend to be concentrated in poorer countries [33], there was no clear pattern between trends of BII and GDP over time, and only a tenuous relationship between temporal trends in BII and GDP per capita in 2001. It is possible that the effect of GDP changes on BII may be masked in our data because of limitations in the treatment of plantation forest. Higher GDP can lead to lower biodiversity is through increased investment into cropland and thus increased rates of forest conversion to plantations [34]; if the impact of plantation forests are not included in projections of biodiversity change, correlations with GDP may be muted.

We have so far only considered the impacts of land-use change and related pressures on biodiversity. Although these are the most important drivers of biodiversity loss in the recent past and near future, particularly in the tropics [56], climate change is likely to become increasingly important over longer timescales [57], especially as forest conversion already leads to strong changes in local temperature, potentially exacerbating future impacts of climatic change [58]. Incorporating both land-use change and climate change impacts could improve estimates of biodiversity change [57, 59].

The average BII values reported here are substantially lower than the recently published global average, where BII across the terrestrial surface was found to be approximately 84.6%[17]. Several factors contribute to this difference. First, the new land-use maps have stricter bounds on the extent of primary forest. Second, we have fitted a model using data from tropical and subtropical forest biomes only, appreciating that there are often regional differences in response to human impacts, both in terms of alpha [20, 23] and beta diversity [22]. Third, we compare biodiversity to a baseline of minimally-used primary vegetation rather than to all primary vegetation as in [17]; in the PREDICTS database, this is the closest available proxy to the ideal ‘pristine’ baseline [15]. In older transformed landscapes, this may simply be the ‘least disturbed’ baseline [60], as these sites may have suffered human impacts either directly or as spillover effects from impacts in the surrounding landscape, masking the true impact of land-use change. In tropical and subtropical forests, however, where many land-use changes are relatively recent, minimally-used primary vegetation may be a more appropriate proxy. These sites have still likely suffered some impacts: truly undisturbed sites are relatively rare and becoming more so [53–55]. Fourth, our models of compositional similarity make more efficient use of data, allowing us to account for additional habitat degradation related to roads and human population density [61]; and use logit rather than log transformation, to better reflect the distribution. Fifth, we have attempted to incorporate into projections the impact of plantation forests (by assuming that plantation forest is most likely to be incorporated into lightly- and intensively-used secondary forest in the land-use maps), which can drive major biodiversity loss, particularly when expanding at the expense of primary forest [20, 23, 62]. Excluding the impact of plantation forests [as in 17] produces slightly higher values (globally, BII is estimated as 64% in 2001 and 62.1% in 2012). New land-use maps that are being produced with finer land-use classes [e.g. 25 in prep.], once downscaled, should also allow more accurate estimates of BII.

We have provided a method for estimating annual change in BII in the hope that this global indicator of local terrestrial biodiversity can better inform policy at national and international levels by highlighting key areas for conservation or restoration and monitoring progress towards conservation and restoration targets. However, there are a number of caveats that must be considered as well as limits to the interpretation of results. Firstly, pressure data vary in their resolution and in their accuracy.

This is particularly important for road networks, which can grow rapidly; however, the data are only available as a static layer, and the completeness varies regionally. Static road layers may still provide insights into biodiversity responses: for instance, roads built pre-2000 were associated with forest loss in the following decade in Borneo [63]. However, other linear infrastructure, such as gas lines, can also have important consequences for biodiversity that are not included here [64]. Human population density was interpolated between time steps and, as the data are downscaled using an areal-weighting approach, its resolution at the pixel level varies depending on the size of the input areal unit [65]. Our models also assume that the response of biodiversity to land-use change is immediate; lagged responses are likely to be common and may be complex [15, 66, 67]. These difficulties mean that, while we have estimated BII at a 30” resolution, the estimates are most suitable for assessing average changes across larger pixels or areas (e.g., at a country level) and across broader time steps, rather than focussing on pixel-by-pixel and year-on-year changes. In future, parameter uncertainty in both the abundance and compositional similarity models could be incorporated to provide uncertainty bounds on estimates and trends. This will be important especially for urban areas, where the data used here are limited and so projected diversity is more uncertain.

The implementation of BII used here may still underestimate biodiversity loss. One reason is that the compositional similarity metric implied by the original definition of BII [16] is quite permissive, in that the species abundance distribution could be completely reorganised without reducing BII, provided that the total abundance of originally-present species is not reduced and novel species are not introduced. Therefore, a region with high BII can still under some circumstances have shown strong losses in other aspects of diversity [See 68 for more details]. Using a combination of beta-diversity metrics may provide a more comprehensive assessment of the state of diversity [47, 48] and is a natural avenue for future development of BII.

Although estimating how BII has changed across space and time includes many underlying assumptions and uncertainty, the approach we have used goes beyond our previous implementation of BII by producing annual estimates based on improved statistical modelling and time-varying data on pressures. These annual estimates provide a useful tool for policy makers hoping to track progress towards national and international targets, and for assessing the state of nature; they also provide further evidence of the perilous state of tropical forest biodiversity [69].

## Methods

### Statistical models of how biodiversity responds to pressures

#### Biodiversity data

The models we use are adapted from those in [17] and are based on the PREDICTS database, a global collation of spatial comparisons [19]. The database contains surveys (‘studies’) of multiple sites differing in land use and related pressures [see 18 for details]. The data were subset to only those sites in the following tropical or subtropical forested biomes: tropical and subtropical coniferous forests, tropical and subtropical dry broadleaf forests and tropical and subtropical moist broadleaf forests [70]. We only used studies where communities were sampled, rather than studies that focused on a single target species. All sites included had known geographic coordinates (so that the geographic distance among sites in a study could be calculated). Where enough information was provided in the methods of the original paper, for each site the sampling grain was estimated as the ‘maximum linear extent’; for example, the total transect length walked when sampling at a site [see 18 for more detail]. The final dataset used for analyses contained 777,173 records from 180 published sources on the abundance of 20,740 species from 5159 sites worldwide (representing 45 countries). Invertebrates make up 42.9% of the species, plants 36%, vertebrates 18.7% and fungi 2.4%.

#### Pressure data

The PREDICTS database holds site-level data on land use (primary vegetation, secondary vegetation, plantation forest, cropland, pasture or urban) and land-use intensity (minimal, light and intense), classified using information in the original sources or provided by their authors [see 18 for details]. Although plantation forest exists as a separate land-use class in the PREDICTS database and is characterised by assemblages that are both relatively low in species richness and compositionally distinct [23, 71], it is rarely separated from other forests in global land-use layers; this is also true here. One option [used by 17] is to model responses to plantation forest but omit the effect when projecting results across space. Given the importance of plantation forests in tropical forested areas, we chose instead to group plantation forests together with secondary vegetation when modelling. We did this because it is the most likely source of plantation forest in the global land-use layers and because previous pan-tropical analyses have shown little difference between losses caused by primary conversion to secondary vegetation or plantation forest [20], unless the plantations are intensive [23]. Lightly- and intensively-used plantation were included with intensively-used secondary vegetation, and minimally-used plantation was included with lightly-used secondary vegetation. In addition to land-use and intensity, we included as pressures human population density [for the year 2005 from 72] and the density of roads [derived from 73]; we estimated these at two spatial scales, using their values within both the 1-km and 50-km grid cells containing each site. Environmental conditions for each site were extracted from WorldClim (elevation, maximum temperature of the warmest month, minimum temperature of the coldest month, precipitation of the wettest and driest month; [74]) so that their effects on beta-diversity across sites could be controlled for.

#### Mixed-effects models

Two mixed-effects models [*lme4* package version 1.1-15, 75] were run. The first model focussed on total abundance, calculated as the sum of abundance across all species recorded at each site. If sampling effort varied among sites within a study, abundance was rescaled assuming that diversity increased linearly with sampling effort. Within each study, total abundance was then rescaled so that the maximum value was unity; this rescaling reduces the inter-study variance caused by differences in sampling effort and taxonomic focus and so facilitates modelling. Rescaled total abundance was square-root transformed prior to modelling, which used Gaussian errors; non-integer abundances in the original data precluded modelling of untransformed values with Poisson errors, and square-root transformation resulted in a better residual distribution than *ln*-transformation. Rescaled total abundance was modelled as a function of the following fixed effects: site-level land use and intensity (LUI), human population density (*ln*(*x* + 1) transformed), and density of roads at the 1km and 50km scale (cube-root transformed), along with two-way interactions of LUI with each other pressure. We included an additional control variable to account for among-study differences in human population density (by taking the mean value within each study); this was to control for potential sampling and detection biases where sampling may be more complete in areas of higher human population density (which are generally closer to research institutions and more accessible for sampling). All continuous variables were centered and scaled to reduce collinearity. We used a random-effect structure of spatial block within study, to account for differences in sampling methodology and large-scale environmental differences across studies and the spatial structure of sites within studies. With the model fitted using Restricted Maximum Likelihood (REML), we assessed whether random slopes were required by comparing Akaike’s Information Criterion (AIC) for models with each variable fitted as a random slope in turn. The best fixed-effects structure was then determined using backwards stepwise model simplification with the model fit using Maximum Likelihood [76].

The second model assessed the response of compositional similarity to human impacts. We excluded studies where sampling effort varied among sites. For studies with at least one site classed as minimally-used primary vegetation (the baseline site), we calculated for each study in turn the compositional similarity of each site to each baseline site, measured as the proportion of site *j*’s individuals that belong to species also present in site *i* (where site *i* is in minimally-used primary vegetation, i.e., an asymmetric version of the abundance-based Jaccard similarity index: [77]). Compositional similarity was *logit* transformed (*car* package, version 2.1-6, [78]; an adjustment of 0.01 was used to account for values of 0 and 1). Compositional similarity between any pair of sites will be influenced by how much more impacted site *j* is than the baseline site *i*, as well as the absolute level of pressure faced by site *j*. For each continuous pressure variable, we therefore include in the models both the value at site *j* as well as the difference in value between site *i* and site *j*. We included geographic distance (*ln*-transformed) and environmental distance calculated as Gower’s dissimilarity [79] using the *gower* package in R, [80] (cube-root transformed) between sites to account for decays in compositional similarity with distance [22]. Geographic distance was divided by the median maximum linear extent in the dataset prior to *ln*-transformation. The land-use contrast was included as a fixed effect along with its interactions with the continuous variables. As this dataset is more restricted than that used for abundance (because only studies that sample minimally-used primary vegetation can be used), we were not able to consider effects of use intensity within land uses, other than for primary vegetation (split into minimally-used primary vegetation and a combined class of lightly- and intensively-used primary vegetation) and secondary vegetation. Finally, we included the mean value of human population density within each study as a control variable. We included Study as a random intercept and assessed whether a random slope was supported by using the same framework as before, choosing the random structure with the lowest AIC value among the models that converged successfully. Backwards stepwise model simplification was performed to simplify the fixed effects structure of the model fit using Maximum Likelihood. Traditional significance tests based on likelihood ratios are not accurate here, because the data used are not independent (as each site is compared to multiple other sites within the same study). We therefore used permutations to determine whether a variable could be excluded from the model without significant loss of explanatory power [81]. We permuted the dataset 1000 times by randomly shuffling compositional similarity measurements within each study and refitting both the full and simplified model with this dataset. We then compared the likelihood ratio of our observed models with the distribution of likelihood ratios from models using the 1000 permuted datasets to assess whether the ratio was significantly higher than expected based on models with the same differences in parameters. Note that this approach to modelling compositional similarity makes fuller use of the data than that used by [17], which compared independent pairs of sites within studies and averaged coefficients across 100 models fitted to different randomly-chosen sets of pairwise comparisons. Our matrix-based approach uses all relevant site comparisons in the same model, allowing us to estimate more fixed effects, but carries with it the need for permutation tests to assess significance of variables and coefficients [81].

Diversity analyses were performed using R Statistical Software [82] version 3.4.3. Prior to modelling, all explanatory variables were assessed for multicollinearity using Generalized Variance Inflation Factors [83] for each model; all values were below 5, indicating acceptable levels of collinearity. Transformation of explanatory variables were chosen based on improvements to residual distribution.

### Global pressure data and maps of BII for each year

#### Land use

Hoskins et al. [24] statistically downscaled global land-use data for the year 2005 from 0.5-degree resolution [25] to 30 arc-second resolution, estimating the fraction of each pixel in each of the following classes: primary habitat, secondary habitat, cropland, pasture and urban. That approach was extended here, by integrating the static data for 2005 with remotely-sensed time-varying data on land cover and forest change.

The original method described in Hoskins et al [24] uses a combination of Generalised Additive Models (GAMs) and constrained optimisation to produce fine-grained predictions of multiple land-use classes using the best-available spatial data on climate, landform, soil, land cover, human population density and accessibility (at 30 arc second resolution) as inputs. We made several modifications to this method in order to generate our land-use time-series. To improve predictions outside of the fitted parameter space, we performed AIC-based backwards stepwise model selection, to identify the most parsimonious set of predictor variables. We then fitted our downscaling models to the year 2005 coarse-grained Land-Use Harmonisation data [25] and, using time-varying covariates, used these models to predict land-use for the full time-series. Our time-varying covariates were derived from a land-cover product with a yearly temporal resolution [84]. Once our downscaling models were fitted to the 2005 data of this land-cover dataset, we were able to predict land-use change using the remaining years in the time-series.

We maximised the influence of the time-varying covariates in our downscaling models by fitting the GAMs in two stages. Initially the GAMs were fitted to only the time-varying covariates (i.e., annual land-cover datasets), allowing these to explain as much variation in the data as possible. The static covariates were fitted only in a second step, so that they were only able to describe variation not already described by the time-varying covariates. This resulted in models that maximised information coming from the time-varying land-cover data and, as such, reflected the temporal change in the land-cover layers as much as possible in our land-use predictions.

Within tropical and sub-tropical forested regions (defined as Tropical and Subtropical Moist Broadleaf Forests, Tropical and Subtropical Dry Broadleaf Forests and Tropical and Subtropical Coniferous forests in [70]), we further refined our land-use estimates by integrating the Global Forest Changes (GFC) dataset [27] using the following rules. Within a cell, when the predicted proportion of primary habitat was greater than observed by GFC, primary habitat was reduced to match the GFC-observed forest cover. All other land-uses were then scaled proportionally to their predicted values to ensure all constraints were met. When the sum of predicted primary and secondary habitat were less than observed GFC data they were scaled proportionally so that their sum matched the GFC data. The remaining three land-uses were then scaled proportionally to ensure all constraints were met. This provided land-use estimates within forest biomes that were consistent with the observed change in the GFC dataset. Note that this procedure can result in occasional increases in the amount of primary vegetation over time.

#### Human population density

We downloaded human population density data for the years 2000, 2005, 2010 and 2015 from [72] (adjusted to match 2015 revision of UN WPP Country Totals). After *ln*(*x* + 1) transformation (the 1 is added to avoid problems caused by zeros in the data), we interpolated data for intervening years by assuming linear change in the transformed value over time. For example, a cell’s value for 2006 is given by 0.8 * value for 2005 + 0.2 * value for 2010.

#### Density of roads

We used a vector map of the world’s roads [73] to derive maps of road density: for each 30 arc-second cell, road length is calculated within a 1km and 50km radius from the centre point of the cell and expressed as density per 30 arc-second cell (approximately 1*km*^2^) of land (using the arcpy functions of LineLength and FocalStatistics, ArcGIS v10.5). In the absence of any global time-series data of roads, we treated this layer as a static, rather than dynamic, pressure in our projections.

#### Land-use intensity

To estimate land-use intensity for each year, we applied the statistical models of[71] of land-use intensity to each year’s data on land use and human population density. Briefly, Newbold et al [71] reclassified the Global Land Systems dataset [85] into land-use/use-intensity combinations and then modelled how the proportional coverage of each combination within each 0.5 degree grid cell depended on the proportion of the grid cell under that land use, human population density and UN sub-region (and all two- and three-way interactions).

#### Maps of modelled BII for each year

We used each year’s maps of land use, land-use intensity and human population density, along with the (static) maps of road density to drive the two statistical models of how biodiversity responds to pressures. Pressure data were not permitted to exceed the ranges found in the biodiversity dataset (the values were capped), to prevent extrapolation beyond our data. For total abundance, the values were back-transformed (squared) and expressed relative to the baseline of minimally-used Primary vegetation with zero human population and road density. For compositional similarity, values were back-transformed (inverse logit with adjustment) and expressed relative to minimally-used Primary vegetation, with zero human population and road density and zero environmental distance and geographic distance (note that for the latter, this equates to the median sampling grain in the dataset). Control variables (study-level mean human population density and environmental variables) were set to zero. Multiplying these spatial projections together produces BII. We did this for each year between 2001 and 2012. Average BII values for each country, subregion and region were calculated for each year by averaging modelled values across all grid cells intersecting the relevant region’s shape file (as defined for the IPBES assessment, from [86]) after re-projecting to a Behrmann equal-area projection. To assess overall trends across the time period, we calculated the log response ratio of start (year 2001) and final (year 2012) values as *ln*(*bii*_2012_*/bii*_2001_). Wilcoxon signed-rank tests were used to assess average trends across all countries. We also relate these changes to contemporaneous changes in GDP per capita (in current US dollars, [87]) and GDP levels at the start of the time series (log-10 transformed value at 2001). We ran linear mixed effects models [88], including IPBES subregion as a random intercept to account for spatial autocorrelation among neighbouring countries. Simulated model residuals were also tested for spatial autocorrelation. For models including the log response ratio of GDP as an explanatory variable, spatial autocorrelation was still evident in the residuals [89–91], so a gaussian spatial autocorrelation structure was included in the model (nlme package, [92]). Models were run on all countries and on only those where at least 50% of their area was included in the projections.

## Acknowledgements

We are grateful to Jason Tylianakis for statistical advice, to Jörn Scharlemann for advice on the manuscript, and to all past and present members of the PREDICTS team who have assisted in data collation and curation. Our thanks also go to all data contributors to the PREDICTS project. PREDICTS is endorsed by the GEO-BON. This is a contribution to the Imperial College Grand Challenges in Ecosystems and the Environment Initiative. This work was supported by the Prince Albert II of Monaco Foundation, NERC (grant NE/J011193/2 and NE/M014533/1 to AP) and a DIF grant from the Natural History Museum.

## Author contributions statement

A.H. and S.F. developed the downscaled land use procedures and data. A.D.P., K.S.O., L.B., T.N. and A.P. designed the methodology for biodiversity analyses, A.D.P. carried out biodiversity analyses. R.E.G. developed the software for performing biodiversity projections over space and time. A.D.P. carried out the biodiversity projections. A.D.P., A.H. and A.P. wrote the first draft of the manuscript. All authors contributed significantly to revisions of the manuscript.

## Additional information

The authors declare no competing interests. The biodiversity used here are openly available for download from the NHM data portal (data.nhm.ac.uk) along with summary statistics for land use and BII for each country and region. The land-use layers area openly available on the CSIRO data portal.

