## Appendix for "Annual changes in the Biodiversity Intactness Index in tropical and subtropical forest biomes, 2001-2012" (from another version of this preprint: version 2, 2019-03-06)

Supplementary Methods:

*Statistical downscaling procedure*

Here we give description of the original method with detailed descriptions of the additional processes used to produce our time-series of land-use change. Additional detail and background on the original downscaling method can be found in Hoskins et al. (2016).

The downscaling method described in Hoskins et al (2016) uses a combination of Generalised Additive Models (GAMs) and constrained optimisation to produce fine-grained estimates of spatial data from regularly gridded coarse-grained information. The results are the estimated proportions of multiple land-use classes within each fine-grained grid cell where all classes are balanced so that their sum equates to 1. This method extends the single class, general downscaling method developed by Malone et al (2012) by including a constrained optimisation step which allows the balanced downscaling of multiple classes of proportional data.

Initially, GAMs are fitted using a quasibinomial error structure and logistic link function to a set of fine-grained predictor variables. The values from the coarse-grained data to be downscaled are set as the response variable and a separate GAM is used for each class to be downscaled (in our case 5). Once all GAMs are fitted the resulting fitted values are rescaled multiplicatively so that the coarse-grained aggregated proportions are equal to the initial coarse-grained input proportions. The subsequent values are then passed to a constrained optimisation algorithm which, using the associated χ^2^ function, estimates the optimal proportions for each land-use class, ensuring that the values remain between 0 and 1 and that the sum of all values equals 1 while taking into account the standard error of each fitted value. The subsequent balanced values are then passed back into the GAMs and the process is repeated. Once the mean inter-iteration prediction difference from a land-use class minimises to within a specified tolerance, this class is held fixed for subsequent iterations. After all land-use classes have converged on a solution, the fitted values are passed back to the constrained optimisation algorithm for a final time to ensure that all cells balance to 1 and the results are returned.

The method we used to create the current, time-series dataset differed in several ways. The GAMs fitted in each iteration of the analysis were fitted in two stages, rather than one. An initial GAM was fitted to only the time-varying predictor variables in the models. The remaining static variables were then fitted in a subsequent model which was only able act on the residual variation from the first step of the process. This ensured that the final predictions were utilising the maximum amount of information available from the time-varying components within the models.

We previously fitted models to a set of predictor variables that were *a priori* determined to have potential relationships with fine-grained land-use patterns. Prior to downscaling our time-series version of these data, we performed backwards step-wise model selection via AIC to determine the most parsimonious set of time-varying and static predictor variables for each downscaling model. These optimal sets of predictor variables were then used in each downscaling model. Additionally, our original models used principal components of the predictor set. However, to ensure that we were describing change in land-cover correctly, we did not take principal components of the final covariate data.

*Generating yearly predictions of global land-use*

Our aim for the time-series downscaled land-use dataset was to downscale a single time point (2005) via our modified downscaling method. We then used the fitted 2005 models to predict land-use for the remaining years (2001-2004 and 2006-2012) by substituting the 2005 land-cover predictors for values from other years.

With the exception of the land-cover data, our set of original fine-grained (30” resolution) predictor variables remained the same as those used in Hoskins et al. (2016). These consisted of a variety of climate, land-form, substrate and anthropogenic influence layers. Each of these layers was considered as stable over the timeframe of our dataset and so were treated as static variables in our downscaling models.

The time-varying component of our models came from the land-cover data. We used the MODIS based land-cover product (MCD12Q1) developed by Friedl et al (2010). This dataset consists of the yearly (currently 2001-2012) categorisation of XX land-cover classes at a resolution of approx. 500m^2^. Each yearly layer was separated into individual binary layers representing the presence or absence of each land-cover class within a cell. These binary layers were resampled using bilinear interpolation to generate layers at 30” resolution (approx. 1km^2^) of the proportion of each land-cover class within each 30” cell, globally. These 30” resolution layers were then included as the time-varying component within our downscaling models.

Spatial separation of the globe into multiple smaller segments for downscaling followed the methods used in Hoskins et al (2016). Briefly, we separated the globe into the 61 bio-realms described in Olson et al (2001). Each segment had an additional 50 km buffer region around it to ensure that information from neighbouring regions was included in each model. When generating a global mosaic of all 61 downscaling models, the buffered cells were removed and neighbouring models were joins without any spatial smoothing between models. For further description and justifications of our spatial segmentation and aggregation choices refer to Hoskins et al (2016).

*Integration of Global Forest Change datasets*

Once we had generated downscaled land-use predictions we further refined our estimates within forested regions by integrating information from the Global Forest Change (GFC) datasets (Hansen et al 2013). These data provide estimates of forest cover at 30 m resolution, in yearly time steps from 2000 to 2014, across the globe. We generated 1 km resolution aggregate estimates from these data and used them to scale our estimates of land-use within forested biomes (as defined by Olsen 2001) so that proportions of primary forest within a cell could not exceed the proportion of forested land as detected by the GFC data.

We did this by enforcing the following rules on the predictions of land-use. Where the proportion primary habitat was predicted to be greater than the observed proportion of forest within a cell, the primary habitat prediction was reduced to match the proportion of forest within a grid cell. All other land-uses were then scaled proportionally to their original predictions, making sure that the sum of all fives land-uses still equalled 1. Where the sum of predicted primary and secondary habitats was less than the observed proportion of forest within a cell, primary and secondary were scaled proportionally so that their sum equalled the proportion of forest observed. Subsequently, the remaining three land-uses were scaled proportionally to their original predictions, making sure that the sum of all five land-uses equalled 1.

Figure S1: Average BII values for each country against their GDP per capita, for 2001 and 2012 (arrows indicate difference between 2001 values and 2012 values of BII). Note some countries are not presented as GDP per capita was not present in the datasource.

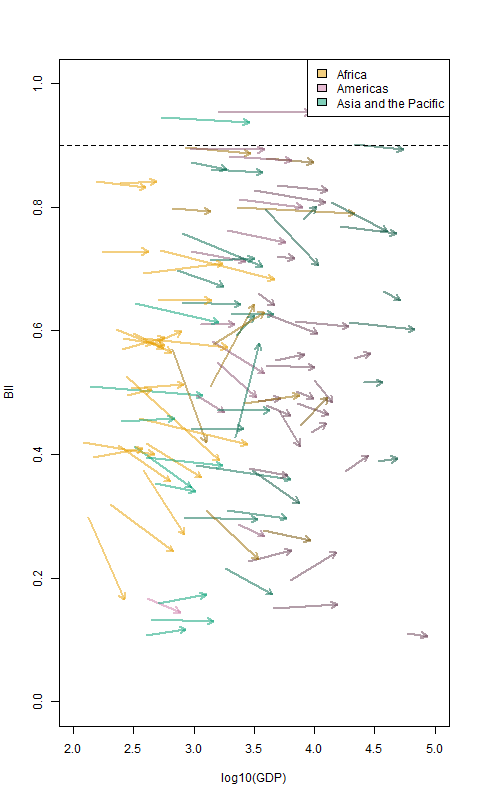
